# Matrix stiffness modulated release of spheroid-derived extracellular vesicles and discovery of Piezo1 cargo

**DOI:** 10.1101/2025.01.13.632826

**Authors:** Maulee Sheth, Manju Sharma, Supasek Kongsomros, Maria Lehn, Takanori Takebe, Vinita Takiar, Trisha Wise-Draper, Somchai Chutipongtanate, Leyla Esfandiari

## Abstract

Augmented extracellular matrix (ECM) stiffness is a mechanical hallmark of cancer. Mechanotransduction studies have extensively probed the mechanisms by which ECM stiffness regulates intracellular communication. However, the influence of stiffness on intercellular communication aiding tumor progression in three-dimensional microenvironments remains unknown. Small extracellular vesicles (EVs) are communicators of altered biophysical cues to distant sites through EV-ECM interactions and EV-mediated recipient cell-ECM interactions. Here we demonstrate stiffness-mediated modulation of small EVs secretion and cargo from three-dimensional oral squamous cell carcinoma spheroids. Using a spheroid culture platform with varying matrix stiffness properties, we show that small EVs carry parental biomolecular cargo, including mechanosensitive Piezo1 ion channel and adhesion molecule CD44. We comprehensively validate the presence of both markers in our EV populations using proteomic and genetic analysis. Transcriptomic analysis of microRNA and long non-coding RNA cargo of small EVs released from soft and stiff ECM spheroids revealed enrichment of tumorigenic and metastatic profiles in EVs from stiff ECM cultures compared to that of soft ones. Gene set enrichment analysis of a comparative dataset obtained by overlaying spheroid mRNA and EV miRNA profiles identified key oncogenic pathways involved in cell-EV crosstalk in the spheroid model.

## Introduction

Extracellular matrix (ECM) stiffening is a key biophysical alteration associated with the progression of multiple solid cancers^1^. Mechanoreceptors, including integrins and mechanosensitive ion channels, sense changes in ECM stiffness, and mechanotransduction of these signals fosters malignant cell behavior^2, 3^. To this end, stiff ECM promotes cell proliferation, epithelial-to-mesenchymal transition, metastasis, and therapeutic resistance^4–6^. While the direct influences of ECM stiffening on tumor cell behavior have been extensively studied in 2D and increasingly in 3D, there is little evidence of the indirect effects employing other modes of communication that promote tumor growth, such as extracellular vesicles (EVs).

Small EVs are 30 to 150 nm membrane-bound nanovesicles released by all cells. EVs released by both healthy and pathological cells are also known to be regulators of the ECM either through direct EV-ECM interactions or by indirectly influencing cell-ECM interactions^7^ (Figure 1). In cancer, small EVs play a crucial role in intercellular communication by facilitating interplay and rewiring between different components of the tumor microenvironment (TME) and contributing to cancer metastasis by enhancing tumor cell migration, remodeling the ECM, and establishing the pre-metastatic niche^8–12^. EVs are rich in their bioactive cargo, including signaling proteins, nucleic acids, and lipids derived from their parental cells that affect the pathophysiology of recipient cells^13^. These unique abilities of EVs to reflect their parental cells of origin and influence recipient cells, make them a unique set of circulating biomarkers in both diagnostic and therapeutic applications.

**Figure 1.**
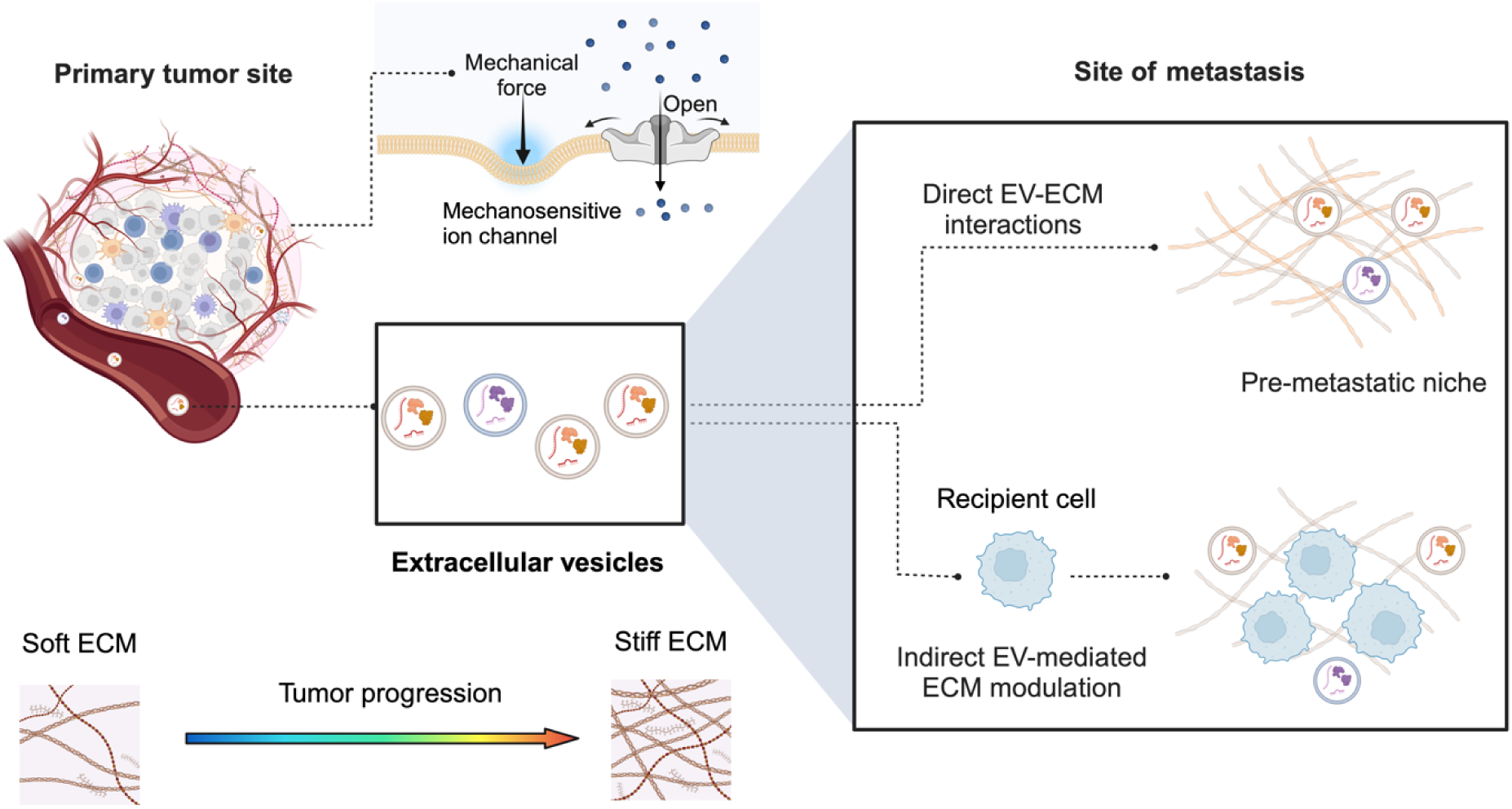
Tissue stiffening is a mechanical hallmark of cancer regulating tumor cell behavior and is perceived by mechanosensitive ion channels. Extracellular vesicles are communicators of these altered biophysical cues to metastatic sites by modulating the ECM directly through EV-ECM interactions or indirectly by influencing recipient cell-ECM interactions.

Recent studies have highlighted the critical role of ECM stiffness in modulating EV behavior, indicating a growing interest in understanding ECM-EV interactions^14, 15^. For example, Sneider *et al.* reported changes in the quantity and protein cargo of primary breast cancer-derived small EVs on stiff ECM, which further aided cancer cell dissemination in a zebrafish xenograft model and induced resident lung fibroblasts to adopt a cancer-associated fibroblast phenotype^16^. Similarly, Liu *et al.* demonstrated that ECM stiffness modulates the secretion and uptake behaviors of mesenchymal stem cell-derived EVs, underscoring the impact of the ECM on EV-mediated intercellular communication^17^. Together, these emerging findings suggest a need for further exploration of ECM stiffness as a determinant of EV-mediated signaling in various pathophysiological contexts. Simultaneously, it has been established that EVs derived from 2D cell cultures dramatically differ from those derived from 3D culture models, as monolayer systems fail to capture the crucial structural and topographical features of 3D *in vivo* microenvironments^18^. In contrast, 3D culture models such as spheroids and organoids have been shown to improve EV yield, enhance therapeutic efficacy, and present more physiologically relevant molecular signatures^19–21^. Despite these advantages, the impact of ECM stiffness on EV dynamics in 3D environments remains largely unexplored. In this study, we address this gap by employing a novel combination approach of employing a three-dimensional culture model to characterize the role of physical tissue properties, i.e., stiffness, on EV secretion and composition from head and neck cancer spheroids.

Head and neck squamous cell carcinoma (HNSCC) is the seventh most common cancer globally^22^. Oral squamous cell carcinoma (OSCC), an HNSCC malignancy, is characterized by increased tissue stiffening which also functions as a biophysical hallmark due to ECM remodeling^23^. Additionally, small EVs are known to have a direct correlation with the aggressiveness of HNSCC^24^. We previously showed matrix stiffness modulates the parental Cal27 OSCC spheroids used in this study^25^. Our results revealed that cancer spheroids respond to static matrix stiffness stimuli through mechanosensory Piezo1 and TRPV4 ion channels and that increased stiffness drives tumorigenic phenotypes in OSCC spheroids. In our current study, we extend the utility of this model to investigate the influence of matrix stiffness on Cal27 spheroid-derived small EVs, including their secretion, ability to reflect parental molecular signatures of interest, and transcriptomic composition.

To our knowledge, this is the first investigation of stiffness-mediated small EVs release in a three-dimensional cancer culture system and the first to show the presence of mechanosensitive Piezo1 ion channels in EVs. We believe it is also the first study to conduct a two-fold RNA sequencing analysis, i.e., miRNA-seq and lncRNA-seq, that advances insight into the impact of matrix stiffness on cancer spheroid-derived EVs composition *in vitro*. Our results indicate that stiff matrix promotes the secretion of small EVs from Cal27 spheroids and that spheroid-derived EVs carry parental biomolecular signatures, including Piezo1 and stemness marker CD44. Lastly, stiff matrix-derived EVs were also found to reveal RNA profiles of relevance to progressed and metastatic cancer.

## Results

### Matrix stiffness promotes the secretion of spheroid-derived EVs

We cultured Cal27, a human oral squamous cell carcinoma (OSCC) cell line, spheroids using the liquid overlay technique on “soft” and “stiff” Matrigel-based environments as per our previously reported study^25^. The elastic moduli of 0.3 kPa and 0.9 kPa for our soft and stiff hydrogels represent the lower range of pathological stiffness of healthy and cancerous oral tissues, respectively. EVs derived from spheroids were isolated from conditioned culture media using an insulator-based dielectrophoretic (iDEP) lab on a chip device and analyzed by nanoparticle tracking analysis (NTA) (Figure 2a). Label-free isolation using the iDEP platform is selective to the 30 to 150 nm size range of small EVs^26–28^. EVs secretion, as marked by particle concentration, increased two-fold for the same number of cells seeded for spheroid formation on the stiff ECM (henceforth termed EV_stiff_) as compared to the soft ECM (henceforth termed EV_soft_) (Figure 2b, c). As such, no significant difference was observed in the average size of EVs between the two groups (Supplemental Figure S1). Additionally, western blot analyses of EV protein markers, including CD63, TSG101, and HSP70, indicated a higher marker abundance for EV_stiff_ with TSG101 being statistically significant (Figure 2d-g).

**Figure 2.**
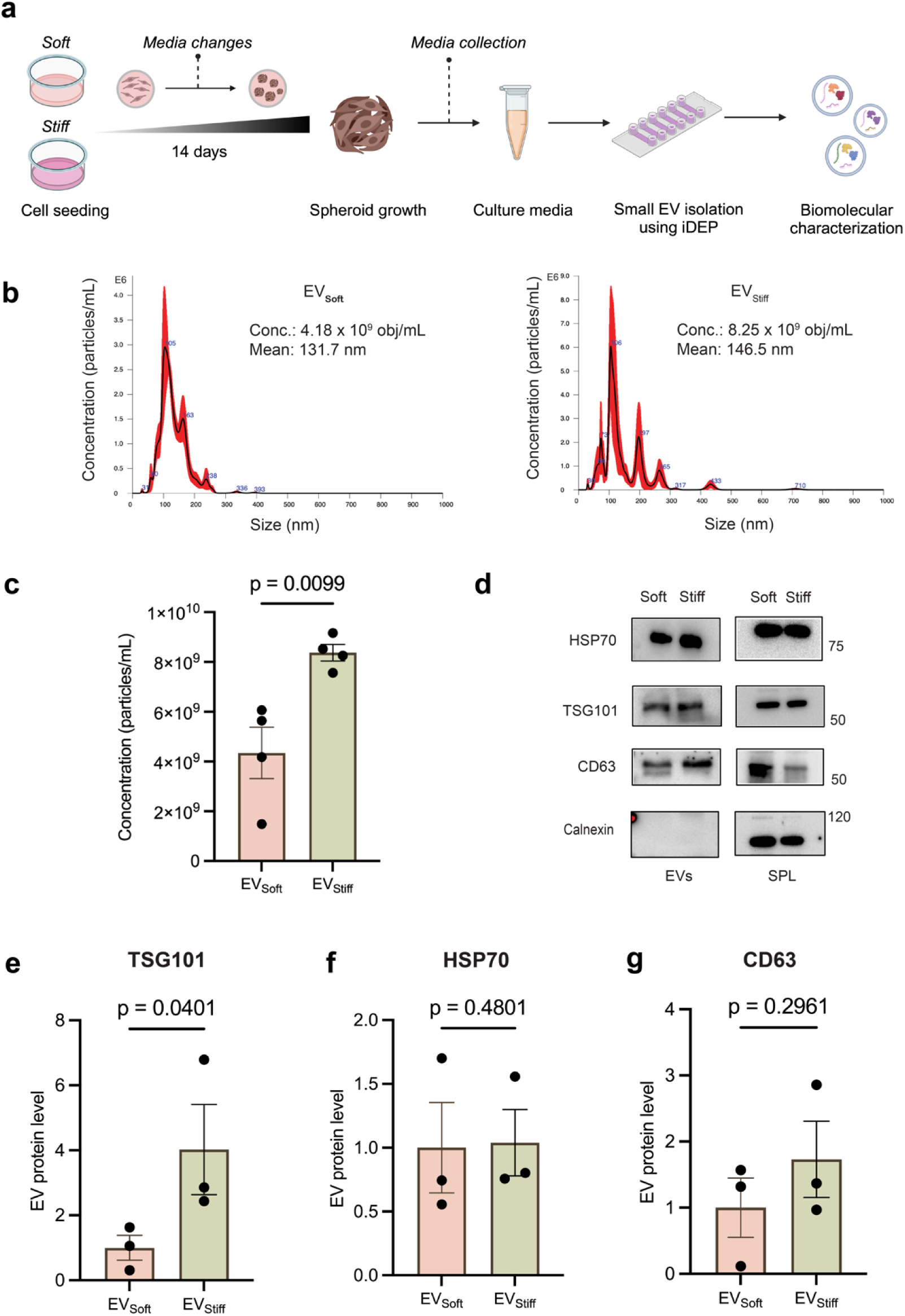
Stiff ECM promotes spheroid-derived small EV secretion. **a)** Workflow indicating experimental timeline and procedure of spheroid culture and small EVs isolation **b)** Representative NTA graphs for purified EV_soft_ and EV_stiff_ samples. **c)** Quantitative analysis of relative nanoparticle secretion between EV_soft_ and EV_stiff_. Values are presented as mean ± SD. n = 4 per group. Unpaired t-test. **d)** Immunoblotting of EV markers (TSG101, HSP70, CD63) in spheroid lysates (SPL) and purified small EVs from Cal27 spheroids cultured in soft and stiff matrices. Calnexin was used as a negative control. Molecular weights (in kDa) are shown to the right. Unprocessed blots are available in the supplementary information file. Quantification of the levels of **e)** TSG101, **f)** HSP70, and **g)** CD63. Proteins for EV_soft_ were normalized as 1. Values are presented as mean ± SD. n = 3 per group. Paired t-test.

### Spheroid-derived EV progenies carry parental Piezo1 and CD44 signatures

We have previously reported the influence of matrix stiffness on mechanosensitive Piezo1 and stemness marker CD44 expressions in Cal27 spheroids with both markers being upregulated in stiff ECM spheroids^25^. Here, we questioned if EVs derived from the culture media of these spheroids carry parental protein markers of interest. Western blot analyses indicated the presence of both, Piezo1 and CD44, in both, EV_soft_ and EV_stiff_ groups (Figure 3a). We further complemented our validation by characterizing the presence of these markers in the EV isolates using imaging flow cytometry (iFCM) and direct stochastic optical reconstruction microscopy (dSTORM). iFCM, with a limit of detection of 70 nm, enabled bulk characterization of Piezo1 and CD44 positivity in EVs along with tetraspanin CD63 (Figure 3b, Supplemental Figure S2). To visualize the presence of Piezo1 and CD44 in individual EVs, colocalization of tetraspanin CD63 with Piezo1/CD44 was assessed in EVs immobilized using phosphatidylserine (PS) through super-resolution microscopy (Figure 3c). The non-protein PS-based capture was also complemented by visualization of Piezo1 and CD44 in EVs immobilized in a protein-based manner using biotinylated CD63 (Supplemental Figure S3). As a final fold of validation, the mRNA presence of Piezo1 and CD44 in EVs was confirmed using next-generation RNA sequencing (Figure 3d, e).

**Figure 3.**
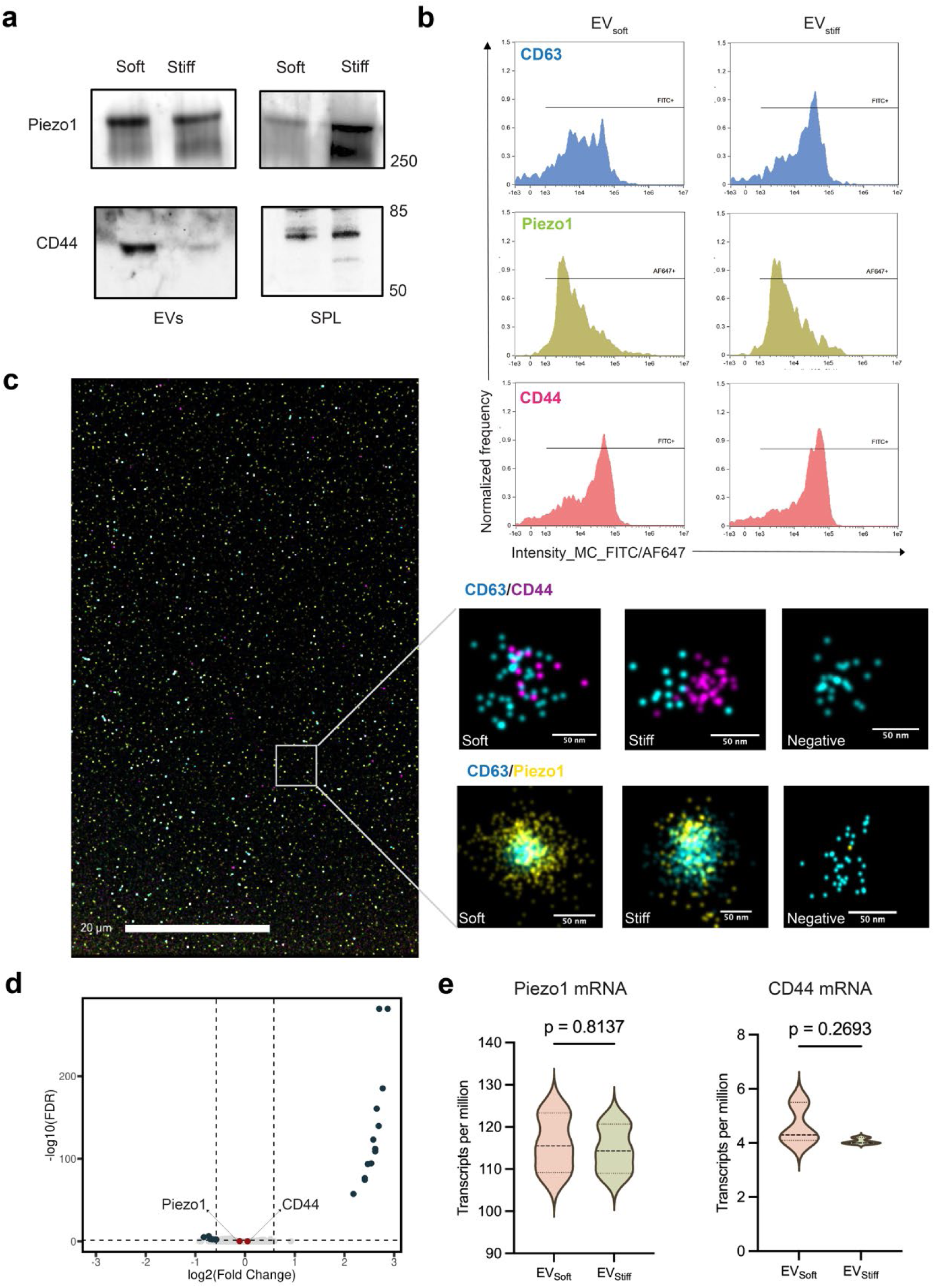
– Spheroid-derived EVs contain CD44 and Piezo1 cargo. **a)** Immunoblotting of new markers (Piezo1, CD44) in spheroid lysates (SPL) and purified small EVs from Cal27 spheroids cultured in soft and stiff matrices. Unprocessed blots are available in the supplementary information file **b)** Imaging flow cytometry for tetraspanin CD63 and new markers Piezo1 and CD44 reveal positive populations in purified EVs from both groups **c)** Representative super-resolution microscopy field of view (left, scale bar = 20 microns) of captured and stained EVs. Single EVs analyzed as double-positive for CD63 and Piezo1/CD44 (right, scale bar = 50 nm) **d)** Volcano plot of differentially expressed mRNAs (log2FoldChange > |0.58| and q-value <= 0.05) indicating the presence of Piezo1 and CD44 mRNAs which are not differentially expressed between the two groups. **e)** Quantitative analysis of Piezo1 and CD44 mRNA presence between the two groups. n=3 per group. Unpaired t-test.

### Matrix stiffness promotes pro-cancerous transcriptome in spheroid-derived EVs

MicroRNAs (miRNA) are vital EV cargo responsible for intercellular and cell-microenvironmental communication in healthy and diseased states. To further investigate how matrix stiffness impacted EV cargo, we performed miRNA sequencing of EV_soft_ and EV_stiff_ spheroid-derived EVs. Principal component analysis indicated distinct clustering of both groups, with 51% and 18% variation explained across PC1 and PC2, respectively (Figure 4a). Differential gene expression analysis revealed 19 miRNAs whose expression levels were significantly modulated with stiffness, as shown in the volcano plot comparing miRNA expression between the groups (Figure 4b). These differentially expressed miRNAs (DEMs) included frequently deregulated miRNAs known to have diagnostic and prognostic roles in head and neck squamous cell carcinoma (HNSCC), such as miR-320, miR-1246, miR-30a, and miR-574 (Figure 4c). Since functions of deregulated miRNAs are highly context-specific with dual roles as either oncogenes or tumor suppressors under certain conditions, we first corroborated our results with previous reports on EV-miRNA trends specific to HNSCC. Oncogenic upregulated DEMs included miR-1246 known to induce movement and invasion of tumor cells in OSCC, miR-30a-3p which regulates Akt signaling to promote metastasis, and miR-455-5p and miR-574-5p prognostic biomarkers^29–32^. miR-122-5p, which is detected solely and universally in small EVs from HNSCC cell lines and promotes metastasis was also upregulated in EV_stiff_. Tumor suppressor downregulated DEMs included miR-200c-3p and miR-125a-5p known to be downregulated in metastatic patients and HNSCC cell line-derived small EVs, respectively^34, 35^.

**Figure 4.**
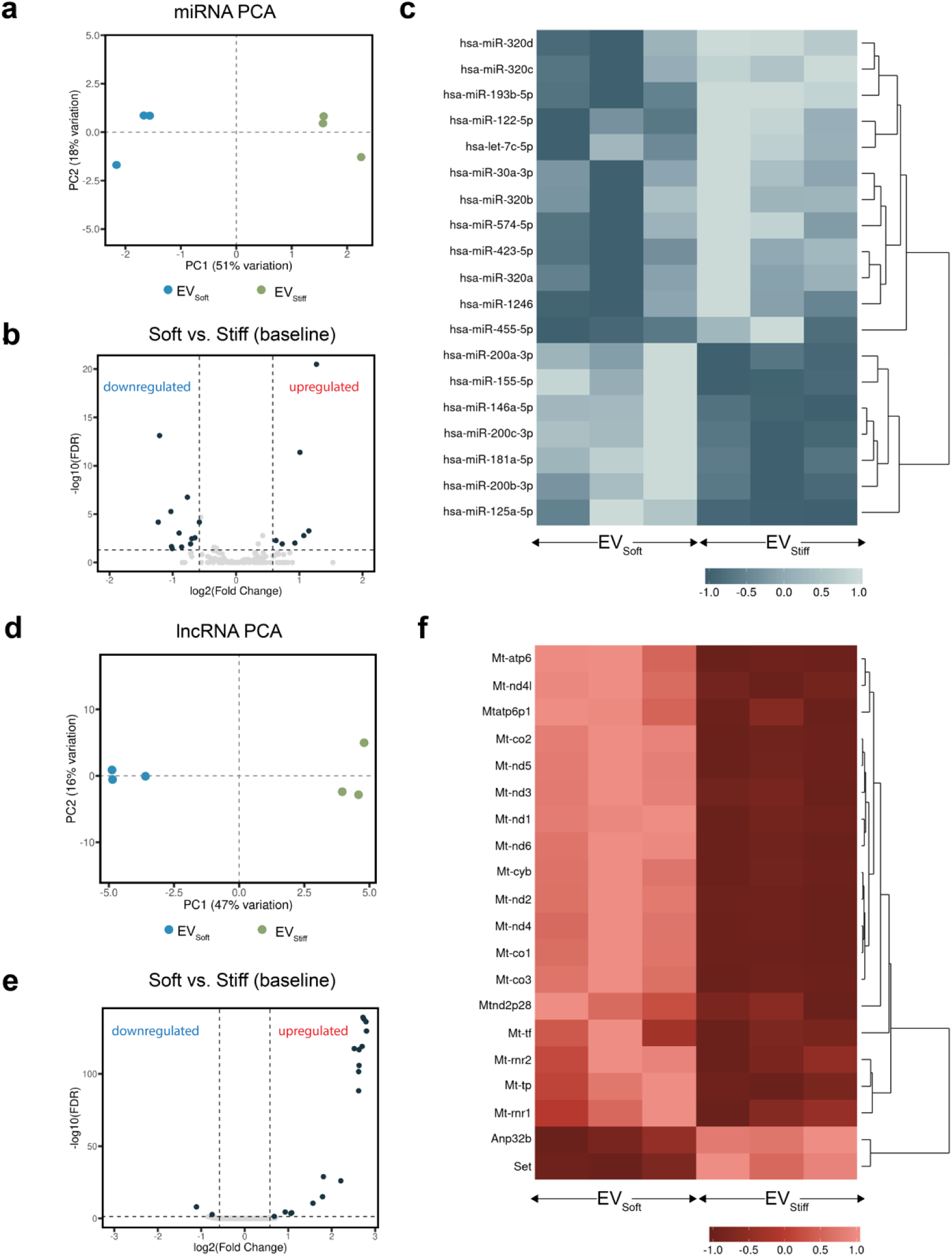
Effect of 3D matrix stiffness on RNA composition of EVs. **a)** Principal component analysis of miRNA-seq wherein PC1 explains 51% of the variance and PC2 explains 18% of the variance. Blue circles indicate EV_soft_ and green circles indicate EV_stiff_ samples (n = 3 biological replicates each) **b)** Volcano plot showing differentially expressed miRNAs (DEMs) (log2FoldChange > |0.58| and q-value <= 0.05) between both groups with EV_stiff_ used as the baseline. Negative fold change indicates relatively increased expression in EV_stiff_ and positive fold change indicates relatively decreased expression in EV_stiff_ quantified by miRNA sequencing **c)** Heatmap showing all 19 DEMs with hierarchical clustering. **d)** Principal component analysis of lncRNA-seq wherein PC1 explains 47% of the variance and PC2 explains 16% of the variance. Blue circles indicate EV_soft_ and green circles indicate EV_stiff_ samples (n = 3 biological replicates) **e)** Volcano plot showing differentially expressed lncRNAs (log2FoldChange > |0.58| and q-value <= 0.05) between both groups with EV_stiff_ used as the baseline. Negative fold change indicates relatively increased expression in EV_stiff_ and positive fold change indicates relatively decreased expression in EV_stiff_ quantified by lncRNA sequencing **f)** Heatmap showing all 20 differentially expressed lncRNAs with hierarchical clustering.

To further investigate how matrix stiffness impacted EV cargo, we investigated the patterns of long non-coding RNAs (lncRNAs) in EV_soft_ and EV_stiff_ spheroid-derived EVs. Principal component analysis indicated distinct clustering of both groups, with 47% variation explained across PC1 and 16% across PC2 (Figure 4d). Differential gene expression analysis revealed 20 lncRNAs whose expression levels were significantly modulated with stiffness, as shown in the volcano plot comparing lncRNA expression between the groups (Figure 4e). Amongst these, multitask oncogene Set, which plays a role in cancer initiation and progression in OSCC, and Anp32b, known to promote cancer growth by regulating c-Myc signaling, were upregulated in EV_stiff_^36, 37^. Interestingly, a majority of the differentially expressed lncRNAs were mitochondrial lncRNAs that were downregulated in EV_stiff_ (Figure 4f). As such, dysregulation of mitochondria-encoded genes has been known to contribute to cancer progression.

### Overlay of spheroid and EV transcriptomic profiles reveal enriched pathways

We next looked at the target genes of EV-DEMs that were identified to be differentially expressed in our previous bulk RNA-seq of parental spheroids to probe cell-EVs crosstalk in our samples (Figure 5a)^38^. Of the 693 target genes of downregulated EV miRNAs, 120 corresponded to upregulated mRNAs in the stiff matrix spheroids (Figure 5b). Conversely, 110 of the 1405 target genes of overexpressed EV miRNAs corresponded to downregulated mRNAs in the stiff matrix spheroids (Figure 5c). We further performed gene set enrichment analysis on the overlapping downregulated DEMs-upregulated DEGs set (Figure 5d). A selection of pathways with pro-cancerous associations, including MAPK cascade, epithelial cell proliferation, and regulation of cell migration and motility, were found to be enriched for the stiff ECM group. Interestingly, certain other enriched pathways included cell communication by electrical coupling, sodium ion transport, and cellular response to endogenous stimuli, suggesting the occurrence of mechano-electrical crosstalk in these microenvironments.

**Figure 5.**
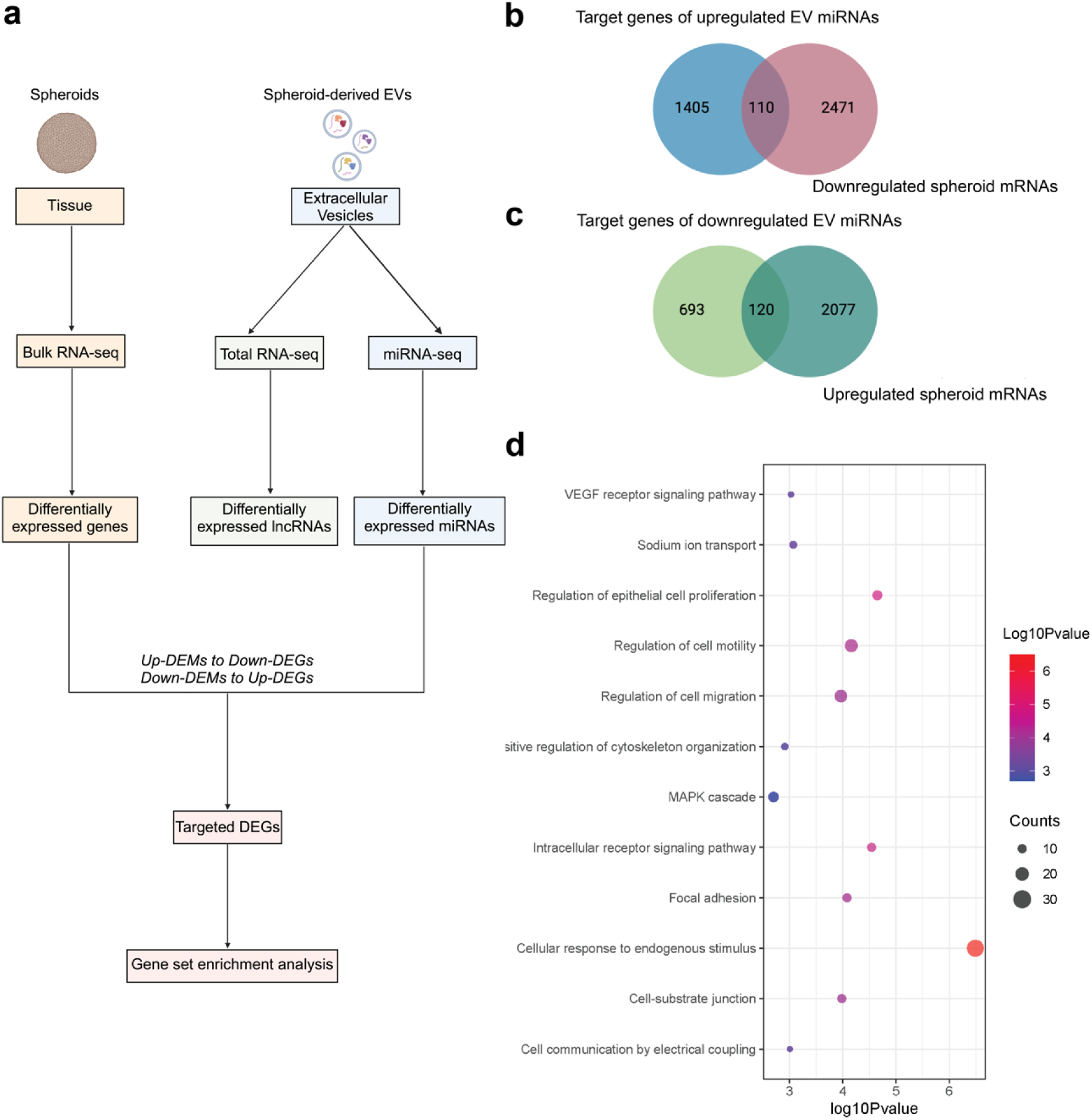
Transcriptomic comparison of spheroidal mRNA and EV miRNAs from soft and stiff ECM cultures. **a)** Workflow indicating miRNA target prediction and data set overlap methodology. Venn diagram showing overlap of **b)** target genes of upregulated EV-DEMs with downregulated spheroid-DEGs and **c)** target genes of downregulated EV-DEMs with upregulated spheroid-DEGs for stiff ECM cultures **d)** Gene set enrichment analysis of the overlapping downregulated DEMs-upregulated DEGs set. The size and color of the circles indicate the number of genes in the dataset associated with the corresponding pathway and the log10Pvalue based on the color bar to the right, respectively.

## Discussion

Our study reports the influence of static biomechanical matrix stiffness stimuli on the secretion and composition of extracellular vesicles released from three-dimensional oral squamous cell carcinoma spheroids. The results reported herein build up on our previous study, where we investigated alterations in Cal27 spheroids in response to “soft” and “stiff” matrix conditions, including the utility of mechanosensitive ion channels in the reception of stiffness cues. Increased ECM stiffness upregulated the secretion of spheroid EVs. The effects of mechanical stimulation on small EV secretion have been previously studied^39^. In particular, our results corroborate previous findings that high matrix stiffness promotes small EV secretion in 2D cell cultures^14, 15^. Wu *et al*. previously revealed a molecular mechanism linking stiff ECM to increased EV secretion by activating intracellular Akt, Rabin8, and Rab8^14^. In another study, Patwardhan *et al.* reported that higher small EVs release on stiff matrices can be attributed to ECM stiffness-dependent activation of the YAP/TAZ pathway^15^. As such, the exact mechanism by which stiffness promotes small EV secretion in our model remains to be investigated.

We have previously investigated expression levels of mechanosensitive ion channel Piezo1 and stemness marker CD44 in Cal27 spheroids cultured in soft and stiff ECM^25^. Probing for the presence of these signatures in progeny EVs led to the discovery of Piezo1 and CD44 in spheroid-derived EV populations from both groups. Enrichment of CD44 cargo in small EVs has previously been reported to promote stiffness-mediated metastasis in breast cancer by facilitating binding to ECM proteins and increasing homing ability to distant organs^16^. The presence of ion channels in small EVs has not been uncovered yet. Our comprehensive characterization revealed the presence of both Piezo1 protein and mRNA in our EV populations. Piezo1 is a mechanosensitive, non-selective cation channel permeable to sodium, potassium, and calcium, and is overexpressed in multiple types of cancerous cells and tissues^40^. The functional significance of EV-Piezo1, if and how EV-Piezo1 levels correlate to cellular-Piezo1 levels under mechanical stimulation, and mechanisms of horizontal transfer of Piezo1 cargo in recipient cells are some interesting avenues warranting further research. As such, calcium signaling, which is a central regulator of tumorigenesis, is known to also regulate EV biogenesis and secretion by modulating Rab GTPase family and membrane fusion factors^41, 42^. While our previous and current studies provide evidence correlating ECM stiffness with Piezo1 activity in cells and with small EV release, respectively, mechanistic insight into how Piezo1-mediated calcium flux and resulting mechanotransduction modulate small EV secretion and composition remains to be investigated.

Given that matrix stiffness had elicited a profound influence on the transcriptomic composition of our spheroids with ∼5000 genes expressed differentially between the two groups as reported in our previous study^25^, we anticipated that RNA sequencing of the small EVs progeny secreted by these spheroids would reveal differential genetic profiles between EV_soft_ and EV_stiff_. We specifically focused on non-coding RNAs including microRNAs and lncRNAs due to their major roles in intercellular communication, ability to impact biological processes in recipient cells, and evident promise as diagnostic and prognostic tools in disease conditions^43–45^. Importantly, since both miRNAs and lncRNAs can have dual roles as both oncogenes and tumor suppressors, we first compared agreement and disagreement in our datasets to EV transcriptomic patterns in head and neck cancers in the literature. In this context, differentially expressed miRNAs (DEMs) upregulated in EV_stiff_ largely correlated to known functional roles as oncogenes. This included miR-1246, miR-122-5p, miR-30a-3p, miR-455-5p, and miR-574-5p known to be upregulated in head and neck cancer EVs and promote invasion and metastasis. Simultaneously, DEMs downregulated in EV_stiff_ correlated to tumor suppressors aiding in cancer progression such as miR-200c-3p and miR-125a-5p, which are known to be downregulated in metastatic patients. Notably, we also saw disagreement in EV-miRNA trends in certain cases, such as miR-200b −3p and −5p, which have been reported to be upregulated in HNSCC cell-derived EVs but were downregulated in EV_stiff_ and let-7c-5p found to be downregulated in HNSCC cell-derived EVs but upregulated in EV_stiff_^33^.

Certain other miRNAs differentially expressed between our groups have not previously been reported in the context of EVs. For these, we broadened our outlook to overall miRNA trends in head and neck cancers. While this allowed us to observe and report the transcriptomic changes in our study, it is important to consider that RNA packaging into EVs is selective, as the RNA profiles in EVs do not fully reflect those in the parental cells. This is a crucial factor when comparing cell- or biofluid-miRNA trends to EV-miRNA patterns. This included upregulated DEMs such as miR-423-5p, miR-193b-5p, and subsets of the miR-320 family, levels of which are upregulated in HNSCC cases with correlation to diagnostic and prognostic significance^46–48^. Finally, downregulated DEMs included miR-146a-5p, miR-155-5p, and miR-181a-5p, which have been negatively correlated with the presence of secondary tumors and distant metastases^48^. Our second fold of sequencing analysis focused on lncRNAs in EV_soft_ and EV_stiff_ isolates. The functional roles and deregulation of EV-lncRNAs in cancer is an incipient research field with limited studies in the literature. To this end, we compared our lncRNA trends to those in head and neck cancers. In this study, we report Set and Anp32b, known to act as oncogenes in cancer initiation and promotion, to be overexpressed in EV_stiff_. Other differentially expressed lncRNAs belong to the mitochondrial family and are generically known to contribute to cancer progression with exact mechanisms warranting further investigation.

In addition to RNA-sequencing of spheroid-derived EVs, our study provides comparative transcriptomic data of spheroidal mRNA and EV miRNAs from soft and stiff ECM cultures^38^. Our previous report on spheroid RNA sequencing identified a selection of genes with associations to mechanosensing, ion channel transport, ECM organization, and tumorigenesis to be upregulated in parental stiff matrix spheroids^25^. In this study, gene ontology analysis of target genes of downregulated EV-miRNAs enriched in the stiff ECM spheroids identified pathways related to tumorigenesis, adhesion, and metastasis. This corroboration is indicative of the substantial role of the miRNA cargo of EVs in cell fate determination of parental cancer spheroids in response to exogenous cues. The enrichment of cell communication by electrical coupling and sodium ion transport pathways in the overlaid dataset, taken together with the upregulation of spheroid genes involved in ion channel transport, also provides evidence of the functional role of EVs in aiding mechanoelectrical crosstalk wherein anomalous biomechanical cues influence the bioelectric state of cancer cells in the TME^49^. Interestingly, an increase in intracellular Na^+^ using ionophores has been reported to cause an initial rise in cytosolic Ca^2+^ and trigger small EVs release by activating the Na^+^/Ca^2+^ exchanger in reverse mode^50^. Since sodium ion channels SCN2B and SCN4B were two of the top five upregulated spheroid genes in the stiff ECM groups, the potential role of ion-channel mediated sodium and calcium ion transport on EV release warrants further investigation.

Traditional two-dimensional cell cultures are the current gold standard to study the biomolecular cargo of EVs secreted from tumor cells for biomarker discovery as well as to investigate the influence of biophysical stimulation on EV secretion and cargo. It has, however, been established that EVs secreted from 2D systems significantly differ in secretion dynamics, molecular signaling contents, and therapeutic effect as compared to those secreted from 3D cultures due to a lack of crucial cell-cell and cell-ECM interactions in 2D^18^. For instance, Thippabhotla et al. demonstrated that 3D culture-derived EV small RNA profiles exhibited a much higher similarity to *in vivo* circulating EVs derived from cervical cancer patient plasma, while 2D culture-derived EV small RNA profiles correlated better with only their parent cells cultured in 2D^20^.

In our study, we utilized a three-dimensional culture platform of cancer spheroids that is more representative of the morphological, structural, and interactional dynamics of *in vivo* tumors. The utility of Matrigel in our platform comes with its set of trade-offs, as others and we have previously discussed^25^. Importantly, since ECM stiffness is generally controlled by varying polymer concentrations, differences in gel microarchitecture are crucial factors influencing particle diffusion. In this regard, Matrigel’s sub-micron effective pore size has been reported to allow for the selective diffusion of only small EVs and soluble factors, preventing the crossing of larger particles such as apoptotic bodies, larger microvesicles, and oncosomes^51^. Considering that EV populations are highly heterogeneous, investigating the influence of matrix stiffness on larger EVs and matrix-bound vesicles purified from harvested hydrogels is an interesting avenue in this regard^52^.

## Conclusion

The crosstalk between altered biophysical cues such as ECM stiffness and small EVs is increasingly being elucidated in the TME. In this study, our findings underscore the role of extracellular vesicles (EVs) as dynamic mediators of the physical TME, demonstrated through the use of a three-dimensional culture system. The identification of two novel molecular cargos, Piezo1 and CD44, in our EV samples presents exciting premise for future studies investigating their diagnostic potential, as these molecules may serve as biomarkers reflecting the mechanobiological properties of the tumor matrix. Furthermore, the ability of EVs to modulate the ECM and influence cellular behavior supports their potential as therapeutic agents. By further elucidating the mechanisms underlying the EV-ECM feedback loop modulating intercellular communication, future research may open new avenues for the development of EV-based diagnostic and therapeutic strategies in cancer. In the long term, this has untapped potential to contribute to the development of EV-based therapies capable of intervening in ECM stiffness-induced oncogenic signaling, cancer progression, and metastasis.

## Materials and Methods

### Cell culture and maintenance

Cal27 cells (ATCC, Manassas, VA) were a kind gift from the Takiar-Wise Draper lab. Cells were cultured in DMEM media (Corning, San Diego, CA) containing 10% fetal bovine serum (Fisher Scientific, Waltham, MA) that was super-depleted of extracellular vesicles via 18-hour ultracentrifugation at 100,000xg^33^, 1% penicillin-streptomycin (Fisher Scientific, Waltham, MA), 1% L-glutamine (Fisher Scientific, Waltham, MA), 1% sodium pyruvate (Fisher Scientific, Waltham, MA), and 1% non-essential amino acids (Fisher Scientific, Waltham, MA).

### 3D cell culture

Cal27 spheroids were generated using the media overlay technique^25^. This method plates cells in media containing a small amount of Matrigel on top of solidified gels to enable cell penetration into the gel for spheroid formation. 24-well plates were coated with a thin layer of 3 or 12 mg/mL Growth factor reduced Matrigel (Corning, San Diego, CA) prepared by dilution in ice-cold PBS and incubated for 30 min to allow the gel to solidify. Batch-to-batch variability of Matrigel was accounted for by utilizing the same batch for replicate experiments. Cal27 cells were trypsinized and counted for seeding. 10,000 cells in 500 μL media containing 1% Matrigel were plated on top of well coatings. Spheroids were cultured at 37 °C and 5% CO_2_ with regular media changes after the first 96 h and every 48 h after that. Day 14 spheroid culture media was stored at −80 °C for purification of EVs.

### Isolation of extracellular vesicles

EVs were isolated from culture media samples using an insulator-based dielectrophoresis approach^26–28^. Culture media samples were centrifuged at 21,000xG at 4 °C for 20 minutes, and the supernatant was utilized for EV extraction. Briefly, glass micropipettes were backfilled with 1X filtered PBS buffer using a 33-gauge Hamilton syringe needle and positioned on a substrate. 50 μL sample and 1X filtered PBS were loaded at the tip side and base side chamber of the micropipette, respectively. EVs were trapped at the tip by applying a 10 V/cm direct current (DC) for 15 min, followed by a release in 15 μL 1X filtered PBS by reversing the applied voltage for another 10 min. 2 mL culture media samples were simultaneously processed by performing parallel aliquots to purify EVs in 600 μL PBS. Purified samples were stored at −80 °C for further analysis.

### Nanoparticle tracking analysis

Purified EVs were diluted in filtered PBS at a dilution of 1:20 and analyzed by nanoparticle tracking analysis (NTA) using a NanoSight NS300 (Malvern, Worcestershire, UK) and the NTA 3.1 software. Camera level 14 and detection threshold 5 were used for instrument settings. Five 60-second recordings were obtained per sample. All post-acquisition functions were at default settings to output the mean, mode, standard deviation, and estimated concentration for each particle size.

### Western blot

EV samples (5 μg protein) were lysed with 1X RIPA buffer and mixed with 4X Leammli buffer (Bio-Rad, Hercules, CA, USA). The samples were heated at 95 °C for 5 min and then run into the NuPAGE™ 12%, Bis-Tris, 1.0 mm, Mini-PROTEAN TGX precast gels for 50 min (Bio-Rad, Hercules, CA, USA) before being transferred onto a PVDF membrane using a turboblot (Bio-Rad, Hercules, CA, USA). The membrane was blocked with Everyblot blocking buffer (Bio-Rad, Hercules, CA, USA) and then probed with 1:1000 anti-CD63 (Abcam, Waltham, MA, USA), anti-HSP70 (Abcam, Waltham, MA, USA), anti-TSG101 (Abcam, Waltham, MA, USA), anti-Piezo1 (Novus Biologicals, Centennial, CO, USA), anti-CD44 (Abcam, Waltham, MA, USA), or anti-Calnexin (Abcam, Waltham, MA, USA) at 4 °C overnight. After washing, the membranes were incubated with 1:2000 goat anti-rabbit IgG secondary antibody HRP (Abcam, Waltham, MA, USA) at room temperature for 1 hour. The immunoblot was developed using Clarity Western ECL Substrate (Bio-Rad, Hercules, CA, USA) and imaged on a ChemiDoc MP Imaging system (Bio-Rad, Hercules, CA, USA). Densitometry for western blotting was quantified using Image J (Fiji). Bands for 8-bit images were selected by a rectangular selection and plotted. Areas were measured by the wand tool and normalized for statistical analyses.

### Super-resolution microscopy

Sub-diffraction limit resolution images of single EVs were obtained using a Nanoimager S Mark II microscope (Oxford Nanoimaging, Oxford, UK) equipped with a 100X, 1.4 NA oil immersion objective, an XYZ closed-loop piezo 736 stage, and dual or triple emission channels split at 640 and 555 nm. ∼1 x 10^9^ EVs in 1 mL were stained with 5 μg/mL mix of antibodies including anti-CD63-Cy38 (Oxford Nanoimaging, Oxford, UK) and anti-Piezo1-FITC (Biolegend, San Diego, CA, USA) conjugated using FITC conjugation kit (Abcam, Waltham, MA, USA) or anti-CD44-APC (Biolegend, San Diego, CA, USA). Antibody mix-only samples were acquired as negative controls. Samples were processed according to the manufacturer’s instructions to immobilize the stained EVs on chips provided with the EV profiler kit. Ten to fifteen fields of view were recorded for each sample using direct stochastical optical reconstruction microscopy (dSTORM). Analysis was performed using algorithms including filtering, drift correction, and DBScan clustering developed by ONI (Oxford Nanoimaging, Oxford, UK) via the Collaborative Discovery (CODI) platform.

### Imaging flow cytometry

Advanced imaging flow cytometry of purified EVs was performed using an ImageStream^X^ Mark II imaging flow cytometer (Amnis, Seattle, WA, USA) at the CCHMC Research Flow Cytometry Core. Fifteen microlitre EV isolates were diluted in 15 μL 1X filtered PBS for flow acquisition. Samples were single stained with 1:20 CD63-FITC (Biolegend, San Diego, CA, USA), CD44-FITC (Biolegend, San Diego, CA, USA), Piezo1-AF647 (Novus Biologicals, Centennial, CO, USA), Isotype IgG1-FITC (Biolegend, San Diego, CA, USA), or Isotype IgG2b-AF647 (Biolegend, San Diego, CA, USA) in the dark for 30 minutes. Unstained, isotype, and antibody-only samples were acquired as negative controls. Data acquisition was performed with low-speed fluidics, high sensitivity, and 7-μm core size at 60X magnification. Channels Ch01 and Ch09 were used for brightfield, and Ch12 was used for side scatter. Data were collected for 3 min/sample for all samples. Analysis was performed using IDEAS software (Amnis, Seattle, WA, USA).

### RNA isolation

Total RNA was extracted from purified EVs using the miRNeasy Micro kit (Qiagen, Valencia, CA) as per the manufacturer’s suggested protocol. Size distribution and estimated RNA concentration were measured using the Agilent 6000 Pico Kit using Bioanalyzer.

### Whole RNA sequencing

Directional whole exosome RNA-seq was performed by the Genomics, Epigenomics and Sequencing Core (GESC) at the University of Cincinnati using established protocols as described previously with updates^53, 54^. To summarize, the quality of total RNA was quality control analyzed by Bioanalyzer (Agilent, Santa Clara, CA). A total of ∼10 ng RNA was input for library preparation using NEBNext Ultra II Directional RNA Library Prep kit (New England BioLabs) under PCR cycle number 9. Library quality control and quantification were performed via Qubit quantification (ThermoFisher, Waltham, MA), and individually indexed libraries were proportionally pooled and sequenced using NextSeq 2000 Sequencer (Illumina, San Diego, CA) under the PE 2×61 bp setting to generate about 44M reads. Upon sequencing completion, fastq files were generated via Illumina BaseSpace Sequence Hub for downstream data analysis.

### Total RNA-sequencing – mRNA analysis

RNA-seq reads in FASTQ format were first subjected to quality control to assess the need for trimming of adapter sequences or bad quality segments. The programs used in these steps were FastQC v0.11.7^55^, Trim Galore! v0.4.2^56^ and cutadapt v1.9.1^57^. The trimmed reads were aligned to the reference human genome version hg38 with the program STAR v2.6.1e^58, 59^. Aligned reads were stripped of duplicate reads with the program sambamba v0.6.8^59^. Gene-level expression was assessed by counting features for each gene, as defined in the NCBI’s RefSeq database^60^. Read counting was done with the program featureCounts v1.6.2 from the Rsubread package^61^. Raw counts were normalized as transcripts per million (TPM). Differential gene expressions between groups of samples were assessed with the R package DESeq2 v1.26.0^62^. Gene list and log2 fold change are used for GSEA analysis^63^ using GO pathway dataset.

### Total RNA-sequencing – lncRNA analysis

RNA-seq reads in FASTQ format were first subjected to quality control to assess the need for trimming of adapter sequences or bad quality segments followed by trimming of adapters. The programs used in these steps were FastQC v0.11.7^55^, Trim Galore! v0.4.2^56^ and cutadapt v1.9.1^57^. The trimmed reads were aligned and quantified using the reference human genome version hg38 from GENCODE.v46^58^ with the program STAR v2.6.1e^59^. Raw counts were normalized as transcripts per million (TPM). Differential gene expressions between groups of samples for the RNA-Seq techniques were assessed with the R package DESeq2 v1.26.0^60^.

### miRNA sequencing

MicroRNA-seq was performed by the GESC at the University of Cincinnati^64, 65^. For library preparation, NEBNext Small RNA Sample Library Preparation kit (NEB, Ipswich, MA) was used with a modified approach for precise miRNA library size selection, which makes it possible for the kit to process low input (ng level) and low-quality (Bioanalyzer RIN value <3) RNA with better library recovery and miRNA reads alignment. Specifically, using ∼8 ng total RNA as input, after 15 cycles of final PCR, libraries with unique indices were first equal-10 µl pooled, column cleaned up, and mixed with a custom-designed DNA ladder that contains 135 and 146 bp purified PCR amplicons. This size range corresponds to a miRNA library with 16-27 nt insert that covers all miRNAs. After high-resolution agarose gel electrophoresis and gel excision, the library pool ranging from 135 to 146 bp, including the DNA marker, was purified and quantified by NEBNext Library Quant kit (NEB) using QuantStudio 5 Real-Time PCR System (Thermofisher, Waltham, MA). The first round of sequencing was performed on NextSeq 2000 sequencer (Illumina, San Diego, CA) to generate a few million reads to quantify the relative concentration of each library. The volume of each library was then adjusted to generate the expected number of equal reads [∼4.5M read clusters] from each sample for final data analysis.

### miRNA-sequencing analysis

The exceRpt pipeline^61^ was utilized for Quality Control of small RNA samples. Gene-level expression was assessed by counting features for each gene, as defined in the NCBI’s RefSeq database^62^. Raw counts were normalized as transcripts per million (TPM).

### Data availability

The data that support the findings of this study are available within the article and its supplementary material.

## Acknowledgments

This study was made possible, in part, using the Cincinnati Children’s Research Flow Cytometry Facility (RFCF; RRID: SCR_022635; supported by NIH Grant No. 1 P30 AR47363) and Informatics Shared Facility (IS4R; RRID: SCR_022622). RNA isolation and sequencing were performed by the UC Genomics, Epigenomics, and Sequencing Core (GESC; supported by CEG grant NIEHS P30-ES006096 to Shuk-Mei Ho). We specifically acknowledge the assistance of Sarah Crosswell (RFCF), Xiang Zhang (GESC), Aditi Paranje (IS4R), Ronika De (IS4R), Ashley Kuenzi (IS4R), and Siobhan King (ONI).

## Funding

This work was funded by the National Science Foundation NSF CAREER ECCS (2046037) to Leyla Esfandiari. Vinita Takiar is supported by the Dr. Bernard S. Aron Endowed Chair.

## Conflict of interest

The authors have no conflicts of interest to declare.

## Ethics approval

Ethics approval not required.

## Author contributions

**Maulee Sheth**: Conceptualization, Methodology, Investigation, Visualization, Writing – original draft, Writing – review & editing, **Manju Sharma**: Methodology, Investigation, **Supasek Kongsomros**: Methodology, Investigation, **Maria Lehn**: Methodology, **Takanori Takebe**: Writing – review & editing, **Vinita Takiar**: Writing – review & editing, **Trisha Wise-Draper**: Writing – review & editing, **Somchai Chutipongtanate**: Investigation, Writing – review & editing, **Leyla Esfandiari**: Conceptualization, Writing – review & editing, Supervision, Funding acquisition.

## Supplementary Information

### A. Relative size of isolated EVs

**Supplementary Figure S1.**
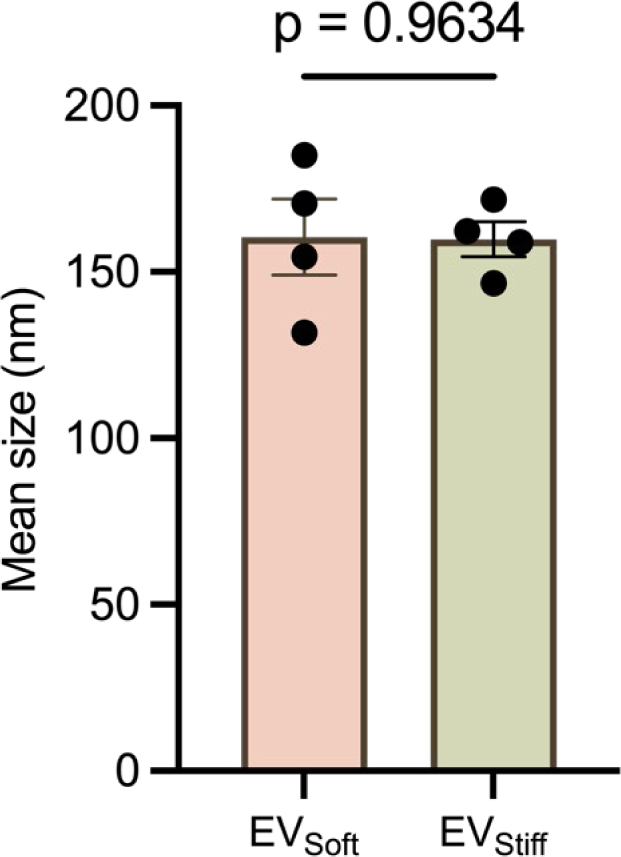
Quantitative analysis of relative nanoparticle size between EV_soft_ and EV_stiff_ measured using NTA. Values are presented as mean ± SD. n = 4 per group. Unpaired t-test.

### B. Imaging flow cytometry controls

**Supplementary Figure S2.**
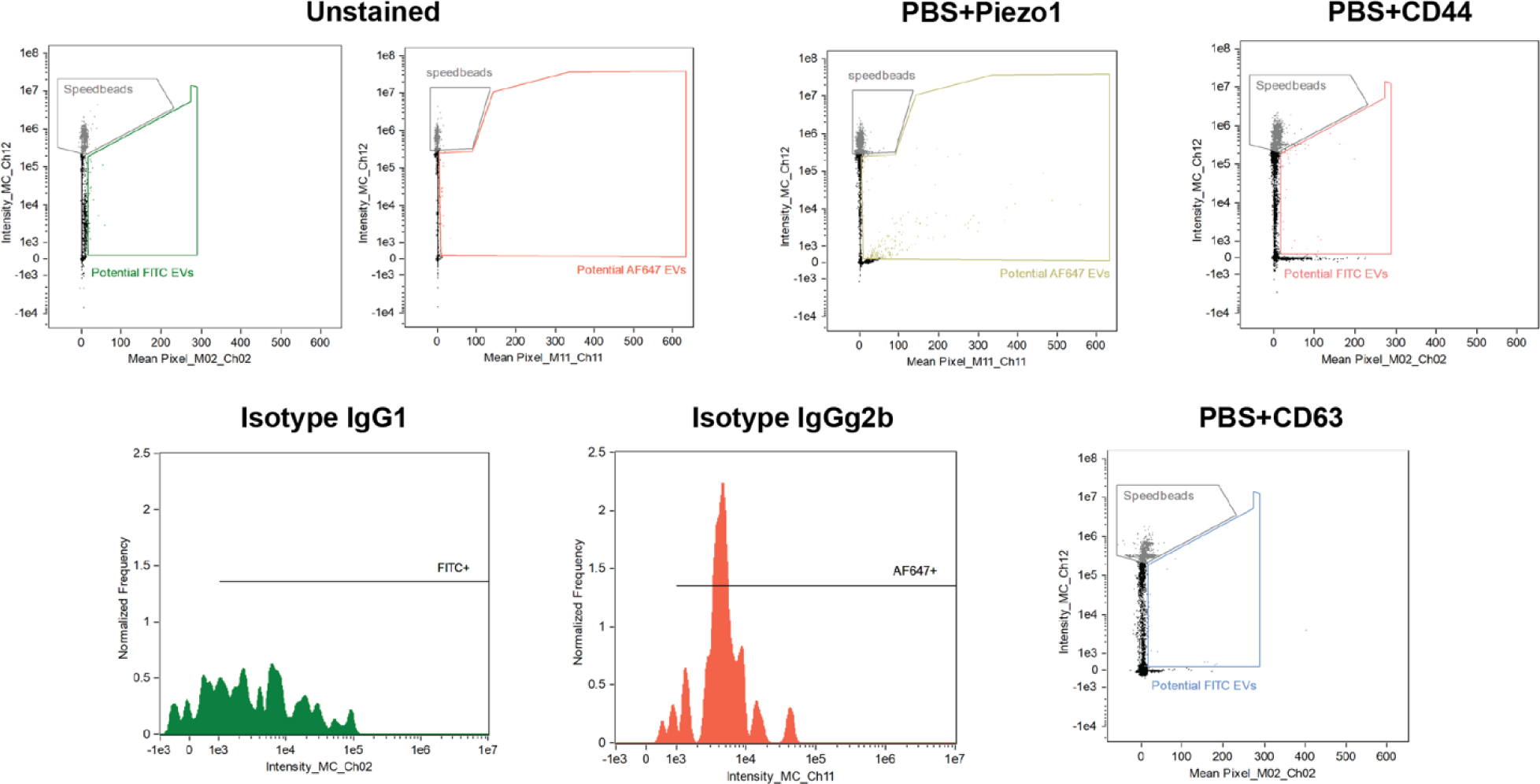
Imaging flow cytometry controls showing gating strategy using unstained samples and negative controls including isotype IgG1 and IgG2b along with PBS + antibody samples for all three markers.

### C. Super resolution microscopy -- biotinylated CD63 capture

**Supplementary Figure S3.**
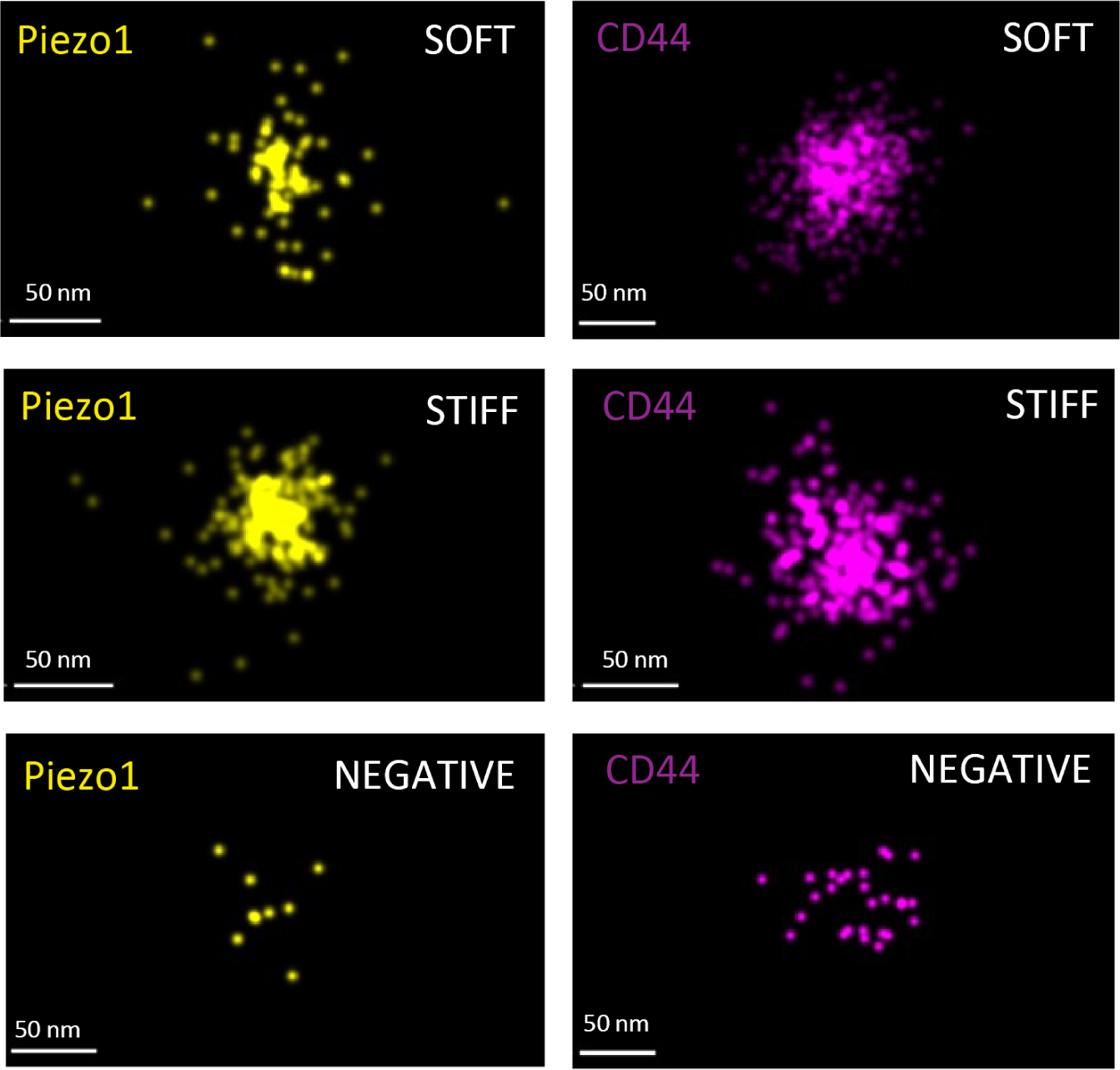
Representative super-resolution microscopy images of single EVs captured using biotinylated CD63 and stained for Piezo1 or CD44 (scale bar = 50 nm).

### D. Immunoblotting raw data

**Supplementary Figure S4.**
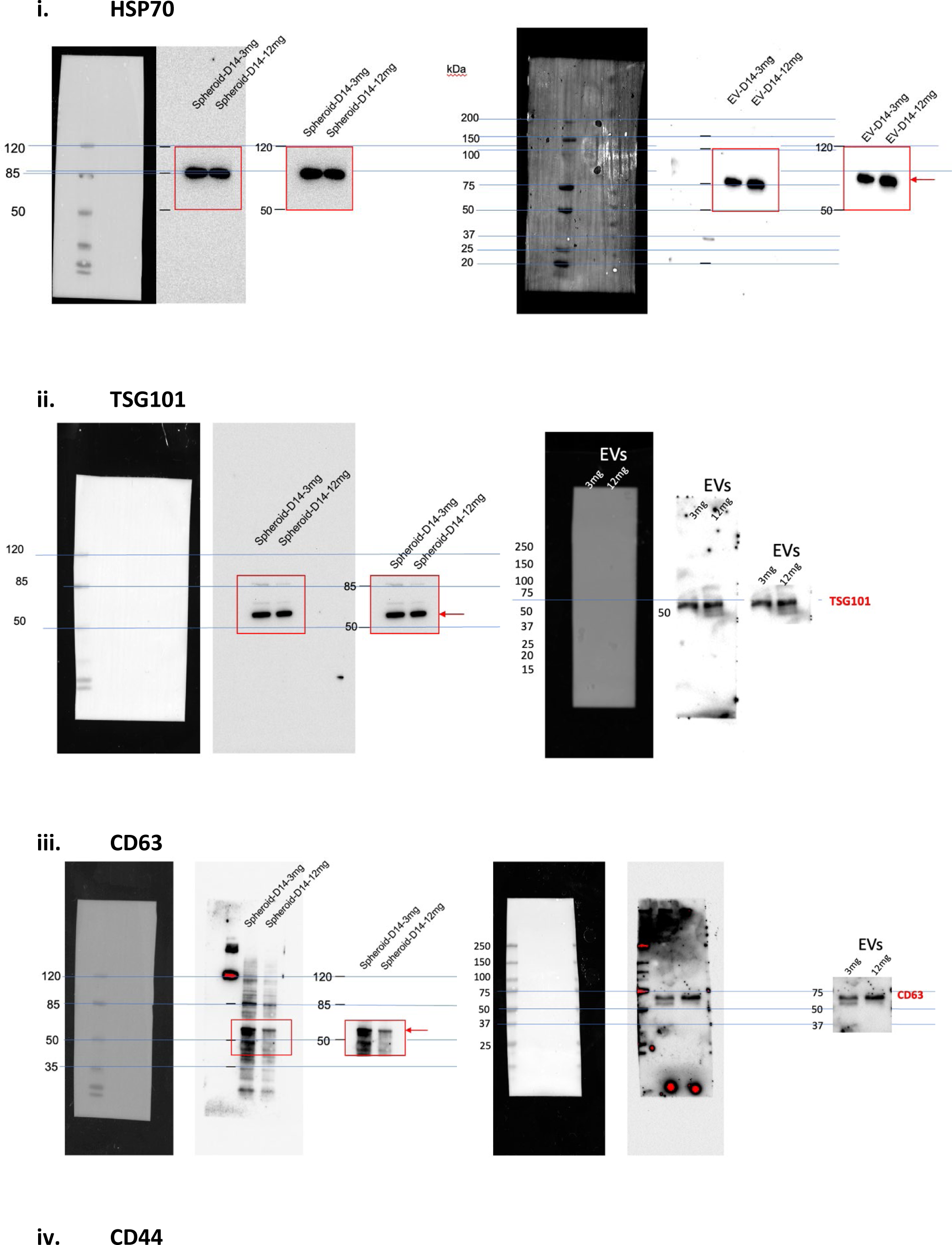
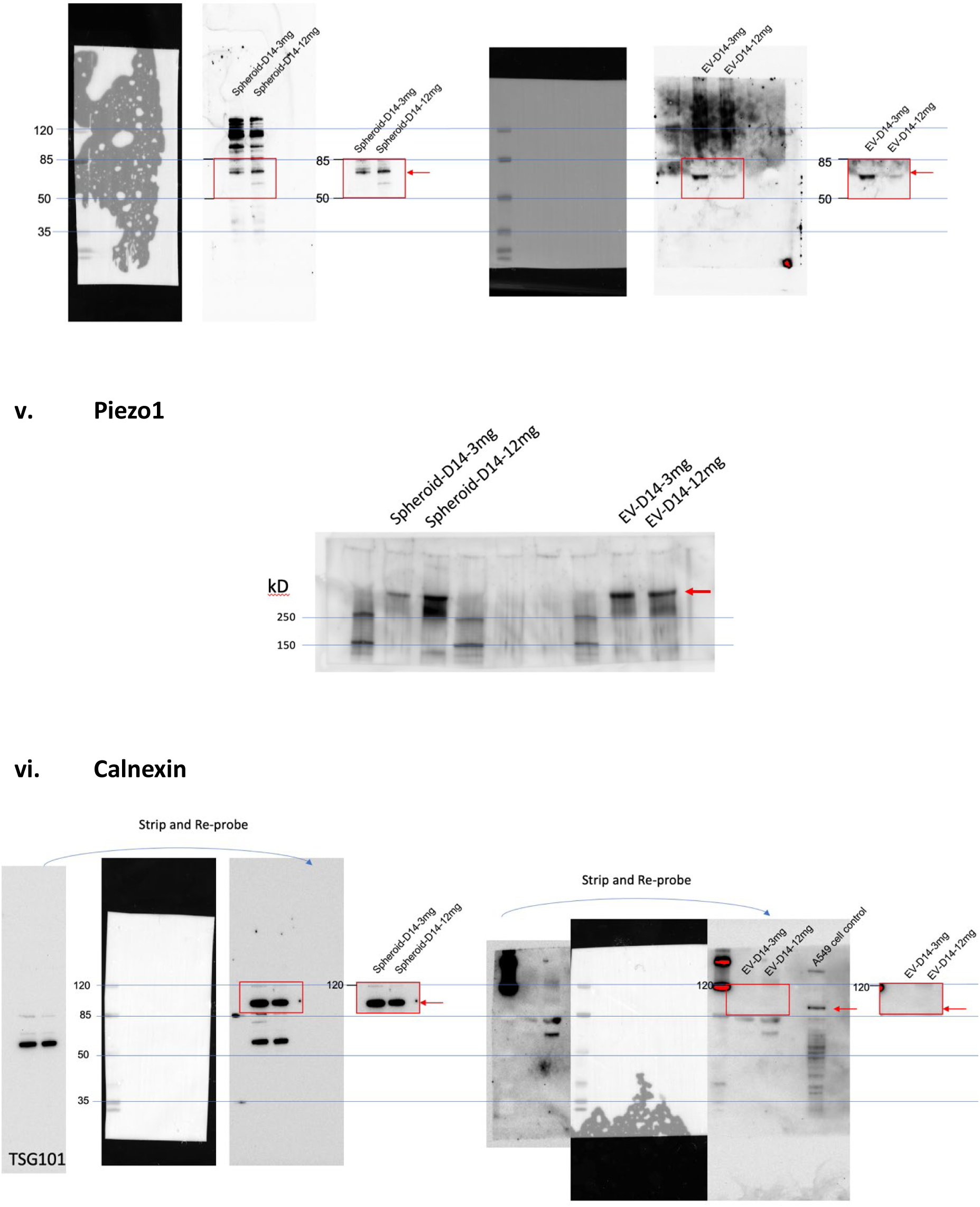
Raw western blot data for EV-specific markers (HSP70, TSG101, CD63), new markers of interest (Piezo1, CD44), and negative EV control (Calnexin) for spheroid lysate and purified EVs samples for both soft (3 mg) and stiff (12 mg) groups.

## Notes

### Competing Interest Statement

The authors have declared no competing interest.

## References

(1) Butcher, D. T.; Alliston, T.; Weaver, V. M. A tense situation: forcing tumour progression. Nature Reviews Cancer 2009, 9 (2), 108–122. DOI: 10.1038/nrc2544.

(2) Jiang, Y.; Zhang, H.; Wang, J.; Liu, Y.; Luo, T.; Hua, H. Targeting extracellular matrix stiffness and mechanotransducers to improve cancer therapy. J Hematol Oncol 2022, 15 (1), 34. DOI: 10.1186/s13045-022-01252-0 From NLM Medline.

(3) Broders-Bondon, F.; Nguyen Ho-Bouldoires, T. H.; Fernandez-Sanchez, M. E.; Farge, E. Mechanotransduction in tumor progression: The dark side of the force. J Cell Biol 2018, 217 (5), 1571–1587. DOI: 10.1083/jcb.201701039 From NLM.

(4) Langley, R. R.; Fidler, I. J. The seed and soil hypothesis revisited--the role of tumor-stroma interactions in metastasis to different organs. Int J Cancer 2011, 128 (11), 2527–2535. DOI: 10.1002/ijc.26031 From NLM Medline.

(5) Li, X.; Wang, J. Mechanical tumor microenvironment and transduction: cytoskeleton mediates cancer cell invasion and metastasis. Int J Biol Sci 2020, 16 (12), 2014–2028. DOI: 10.7150/ijbs.44943 From NLM Medline.

(6) Esfandiari, L.; Paff, M.; Tang, W. C. Initial studies of mechanical compression on neurogenesis with neonatal neural stem cells. Nanomedicine: Nanotechnology, Biology and Medicine 2012, 8 (4), 415–418. DOI: 10.1016/j.nano.2012.01.001.

(7) Patel, N. J.; Ashraf, A.; Chung, E. J. Extracellular Vesicles as Regulators of the Extracellular Matrix. Bioengineering 2023, 10 (2), 136.

(8) Sung, B. H.; Ketova, T.; Hoshino, D.; Zijlstra, A.; Weaver, A. M. Directional cell movement through tissues is controlled by exosome secretion. Nature communications 2015, 6 (1), 7164.

(9) Hoshino, D.; Kirkbride, K. C.; Costello, K.; Clark, E. S.; Sinha, S.; Grega-Larson, N.; Tyska, M. J.; Weaver, A. M. Exosome secretion is enhanced by invadopodia and drives invasive behavior. Cell reports 2013, 5 (5), 1159–1168.

(10) Hood, J. L.; San, R. S.; Wickline, S. A. Exosomes released by melanoma cells prepare sentinel lymph nodes for tumor metastasis. Cancer research 2011, 71 (11), 3792–3801.

(11) Costa-Silva, B.; Aiello, N. M.; Ocean, A. J.; Singh, S.; Zhang, H.; Thakur, B. K.; Becker, A.; Hoshino, A.; Mark, M. T.; Molina, H. Pancreatic cancer exosomes initiate pre-metastatic niche formation in the liver. Nature cell biology 2015, 17 (6), 816–826.

(12) Wortzel, I.; Dror, S.; Kenific, C. M.; Lyden, D. Exosome-mediated metastasis: communication from a distance. Developmental cell 2019, 49 (3), 347–360.

(13) Mathieu, M.; Martin-Jaular, L.; Lavieu, G.; Théry, C. Specificities of secretion and uptake of exosomes and other extracellular vesicles for cell-to-cell communication. Nature cell biology 2019, 21 (1), 9–17.

(14) Wu, B.; Liu, D.-A.; Guan, L.; Myint, P. K.; Chin, L.; Dang, H.; Xu, Y.; Ren, J.; Li, T.; Yu, Z.;, et al. Stiff matrix induces exosome secretion to promote tumour growth. Nature Cell Biology 2023, 25 (3), 415–424. DOI: 10.1038/s41556-023-01092-1.

(15) Patwardhan, S.; Mahadik, P.; Shetty, O.; Sen, S. ECM stiffness-tuned exosomes drive breast cancer motility through thrombospondin-1. Biomaterials 2021, 279, 121185. DOI: 10.1016/j.biomaterials.2021.121185.

(16) Sneider, A.; Liu, Y.; Starich, B.; Du, W.; Nair, P. R.; Marar, C.; Faqih, N.; Ciotti, G. E.; Kim, J. H.; Krishnan, S.;, et al. Small Extracellular Vesicles Promote Stiffness-mediated Metastasis. Cancer Research Communications 2024, 4 (5), 1240–1252. DOI: 10.1158/2767-9764.Crc-23-0431 (acccessed 11/11/2024).

(17) Liu, Z.; Liu, Y.; Li, Y.; Xu, S.; Wang, Y.; Zhu, Y.; Jiang, C.; Wang, K.; Zhang, Y.; Wang, Y. ECM stiffness affects cargo sorting into MSC-EVs to regulate their secretion and uptake behaviors. J Nanobiotechnology 2024, 22 (1), 124. DOI: 10.1186/s12951-024-02411-w From NLM.

(18) Casajuana Ester, M.; Day, R. M. Production and Utility of Extracellular Vesicles with 3D Culture Methods. Pharmaceutics 2023, 15 (2). DOI: 10.3390/pharmaceutics15020663 From NLM.

(19) Cao, J.; Wang, B.; Tang, T.; Lv, L.; Ding, Z.; Li, Z.; Hu, R.; Wei, Q.; Shen, A.; Fu, Y.;, et al. Three-dimensional culture of MSCs produces exosomes with improved yield and enhanced therapeutic efficacy for cisplatin-induced acute kidney injury. Stem Cell Research & Therapy 2020, 11 (1), 206. DOI: 10.1186/s13287-020-01719-2.

(20) Thippabhotla, S.; Zhong, C.; He, M. 3D cell culture stimulates the secretion of in vivo like extracellular vesicles. Scientific Reports 2019, 9 (1), 13012. DOI: 10.1038/s41598-019-49671-3.

(21) Yousafzai, N. A.; El Khalki, L.; Wang, W.; Szpendyk, J.; Sossey-Alaoui, K. Advances in 3D Culture Models to Study Exosomes in Triple-Negative Breast Cancer. Cancers 2024, 16 (5), 883.

(22) Barsouk, A.; Aluru, J. S.; Rawla, P.; Saginala, K.; Barsouk, A. Epidemiology, Risk Factors, and Prevention of Head and Neck Squamous Cell Carcinoma. Med Sci (Basel) 2023, 11 (2). DOI: 10.3390/medsci11020042 From NLM.

(23) Pogoda, K.; Ciesluk, M.; Deptula, P.; Tokajuk, G.; Piktel, E.; Krol, G.; Reszec, J.; Bucki, R. Inhomogeneity of stiffness and density of the extracellular matrix within the leukoplakia of human oral mucosa as potential physicochemical factors leading to carcinogenesis. Transl Oncol 2021, 14 (7), 101105. DOI: 10.1016/j.tranon.2021.101105 From NLM PubMed-not-MEDLINE.

(24) Teng, Y.; Gao, L.; Loveless, R.; Rodrigo, J. P.; Strojan, P.; Willems, S. M.; Nathan, C. A.; Mäkitie, A. A.; Saba, N. F.; Ferlito, A. The Hidden Link of Exosomes to Head and Neck Cancer. Cancers (Basel) 2021, 13 (22). DOI: 10.3390/cancers13225802 From NLM.

(25) Sheth, M.; Sharma, M.; Lehn, M.; Reza, H.; Takebe, T.; Takiar, V.; Wise-Draper, T.; Esfandiari, L. Three-dimensional matrix stiffness modulates mechanosensitive and phenotypic alterations in oral squamous cell carcinoma spheroids. APL Bioeng 2024, 8 (3), 036106. DOI: 10.1063/5.0210134 From NLM PubMed-not-MEDLINE.

(26) Shi, L.; Rana, A.; Esfandiari, L. A low voltage nanopipette dielectrophoretic device for rapid entrapment of nanoparticles and exosomes extracted from plasma of healthy donors. Sci Rep 2018, 8 (1), 6751. DOI: 10.1038/s41598-018-25026-2.

(27) Shi, L.; Kuhnell, D.; Borra, V. J.; Langevin, S. M.; Nakamura, T.; Esfandiari, L. Rapid and label-free isolation of small extracellular vesicles from biofluids utilizing a novel insulator based dielectrophoretic device. Lab Chip 2019, 19 (21), 3726–3734. DOI: 10.1039/c9lc00902g.

(28) Sharma, M.; Sheth, M.; Poling, H. M.; Kuhnell, D.; Langevin, S. M.; Esfandiari, L. Rapid purification and multiparametric characterization of circulating small extracellular vesicles utilizing a label-free lab-on-a-chip device. Scientific Reports 2023, 13 (1), 18293. DOI: 10.1038/s41598-023-45409-4.

(29) Sakha, S.; Muramatsu, T.; Ueda, K.; Inazawa, J. Exosomal microRNA miR-1246 induces cell motility and invasion through the regulation of DENND2D in oral squamous cell carcinoma. Sci Rep 2016, 6, 38750. DOI: 10.1038/srep38750 From NLM.

(30) Chuang, Y.-T.; Tang, J.-Y.; Shiau, J.-P.; Yen, C.-Y.; Chang, F.-R.; Yang, K.-H.; Hou, M.-F.; Farooqi, A. A.; Chang, H.-W. Modulating Effects of Cancer-Derived Exosomal miRNAs and Exosomal Processing by Natural Products. Cancers 2023, 15 (1), 318.

(31) Zhang, H.; Chen, Z.; Huang, Q.; Guo, Y.; Wang, M.; Wu, C. Preliminary study using a small plasma extracellular vesicle miRNA panel as a potential biomarker for early diagnosis and prognosis in laryngeal cancer. Cellular Oncology 2023, 46 (4), 1015–1030. DOI: 10.1007/s13402-023-00792-y.

(32) Yamana, K.; Inoue, J.; Yoshida, R.; Sakata, J.; Nakashima, H.; Arita, H.; Kawaguchi, S.; Gohara, S.; Nagao, Y.; Takeshita, H.;, et al. Extracellular vesicles derived from radioresistant oral squamous cell carcinoma cells contribute to the acquisition of radioresistance via the miR-503-3p-BAK axis. Journal of Extracellular Vesicles 2021, 10 (14), e12169. DOI: 10.1002/jev2.12169.

(33) Langevin, S.; Kuhnell, D.; Parry, T.; Biesiada, J.; Huang, S.; Wise-Draper, T.; Casper, K.; Zhang, X.; Medvedovic, M.; Kasper, S. Comprehensive microRNA-sequencing of exosomes derived from head and neck carcinoma cells in vitro reveals common secretion profiles and potential utility as salivary biomarkers. Oncotarget 2017, 8 (47), 82459–82474. DOI: 10.18632/oncotarget.19614 From NLM.

(34) Galiveti, C. R.; Kuhnell, D.; Biesiada, J.; Zhang, X.; Kelsey, K. T.; Takiar, V.; Tang, A. L.; Wise-Draper, T. M.; Medvedovic, M.; Kasper, S.;, et al. Small extravesicular microRNA in head and neck squamous cell carcinoma and its potential as a liquid biopsy for early detection. Head & Neck 2023, 45 (1), 212–224. DOI: 10.1002/hed.27231.

(35) Dias, F.; Teixeira, A. L.; Nogueira, I.; Morais, M.; Maia, J.; Bodo, C.; Ferreira, M.; Silva, A.; Vilhena, M.; Lobo, J.;, et al. Extracellular Vesicles Enriched in hsa-miR-301a-3p and hsa-miR-1293 Dynamics in Clear Cell Renal Cell Carcinoma Patients: Potential Biomarkers of Metastatic Disease. Cancers (Basel) 2020, 12 (6). DOI: 10.3390/cancers12061450 From NLM.

(36) Zhou, C.; Ma, H.; Yu, W.; Zhou, Y.; Zhang, X.; Meng, Y.; Chen, C.; Zhang, J.; Shi, G. ANP32B inhibition suppresses the growth of prostate cancer cells by regulating c-Myc signaling. Biochemical and Biophysical Research Communications 2024, 698, 149543. DOI: 10.1016/j.bbrc.2024.149543.

(37) Sousa, L. O.; Sobral, L. M.; de Almeida, L. O.; Garcia, C. B.; Greene, L. J.; Leopoldino, A. M. SET protein modulates H4 histone methylation status and regulates miR-137 level in oral squamous cell carcinoma. Epigenomics 2020, 12 (6), 475–485. DOI: 10.2217/epi-2019-0181 From NLM.

(38) Sanchez, I. I.; Nguyen, T. B.; England, W. E.; Lim, R. G.; Vu, A. Q.; Miramontes, R.; Byrne, L. M.; Markmiller, S.; Lau, A. L.; Orellana, I.;, et al. Huntington’s disease mice and human brain tissue exhibit increased G3BP1 granules and TDP43 mislocalization. The Journal of Clinical Investigation 2021, 131 (12). DOI: 10.1172/JCI140723.

(39) Luo, L.; Wu, Z.; Wang, Y.; Li, H. Regulating the production and biological function of small extracellular vesicles: current strategies, applications and prospects. Journal of Nanobiotechnology 2021, 19 (1), 422. DOI: 10.1186/s12951-021-01171-1.

(40) De Felice, D.; Alaimo, A. Mechanosensitive Piezo Channels in Cancer: Focus on altered Calcium Signaling in Cancer Cells and in Tumor Progression. Cancers (Basel) 2020, 12 (7). DOI: 10.3390/cancers12071780 From NLM PubMed-not-MEDLINE.

(41) Alam, M. R.; Rahman, M. M.; Li, Z. The link between intracellular calcium signaling and exosomal PD-L1 in cancer progression and immunotherapy. Genes Dis 2024, 11 (1), 321–334. DOI: 10.1016/j.gendis.2023.01.026 From NLM.

(42) Ambattu, L. A.; Ramesan, S.; Dekiwadia, C.; Hanssen, E.; Li, H.; Yeo, L. Y. High frequency acoustic cell stimulation promotes exosome generation regulated by a calcium-dependent mechanism. Communications Biology 2020, 3 (1), 553. DOI: 10.1038/s42003-020-01277-6.

(43) Sun, Z.; Yang, S.; Zhou, Q.; Wang, G.; Song, J.; Li, Z.; Zhang, Z.; Xu, J.; Xia, K.; Chang, Y.;, et al. Emerging role of exosome-derived long non-coding RNAs in tumor microenvironment. Molecular Cancer 2018, 17 (1), 82. DOI: 10.1186/s12943-018-0831-z.

(44) Dragomir, M.; Chen, B.; Calin, G. A. Exosomal lncRNAs as new players in cell-to-cell communication. Transl Cancer Res 2018, 7 (Suppl 2), S243–s252. DOI: 10.21037/tcr.2017.10.46 From NLM.

(45) Li, C.; Ni, Y.-Q.; Xu, H.; Xiang, Q.-Y.; Zhao, Y.; Zhan, J.-K.; He, J.-Y.; Li, S.; Liu, Y.-S. Roles and mechanisms of exosomal non-coding RNAs in human health and diseases. Signal Transduction and Targeted Therapy 2021, 6 (1), 383. DOI: 10.1038/s41392-021-00779-x.

(46) Romani, C.; Salviato, E.; Paderno, A.; Zanotti, L.; Ravaggi, A.; Deganello, A.; Berretti, G.; Gualtieri, T.; Marchini, S.; D’Incalci, M.;, et al. Genome-wide study of salivary miRNAs identifies miR-423-5p as promising diagnostic and prognostic biomarker in oral squamous cell carcinoma. Theranostics 2021, 11 (6), 2987–2999. DOI: 10.7150/thno.45157 From NLM.

(47) Victoria Martinez, B.; Dhahbi, J. M.; Nunez Lopez, Y. O.; Lamperska, K.; Golusinski, P.; Luczewski, L.; Kolenda, T.; Atamna, H.; Spindler, S. R.; Golusinski, W.;, et al. Circulating small non-coding RNA signature in head and neck squamous cell carcinoma. Oncotarget 2015, 6 (22), 19246–19263. DOI: 10.18632/oncotarget.4266 From NLM.

(48) Thomaidou, A. C.; Batsaki, P.; Adamaki, M.; Goulielmaki, M.; Baxevanis, C. N.; Zoumpourlis, V.; Fortis, S. P. Promising Biomarkers in Head and Neck Cancer: The Most Clinically Important miRNAs. International Journal of Molecular Sciences 2022, 23 (15), 8257.

(49) Sheth, M.; Esfandiari, L. Bioelectric Dysregulation in Cancer Initiation, Promotion, and Progression. Front Oncol 2022, 12, 846917. DOI: 10.3389/fonc.2022.846917 From NLM PubMed-not-MEDLINE.

(50) Savina, A.; Furlán, M.; Vidal, M.; Colombo, M. I. Exosome Release Is Regulated by a Calcium-dependent Mechanism in K562 Cells*. Journal of Biological Chemistry 2003, 278 (22), 20083–20090. DOI: 10.1074/jbc.M301642200.

(51) Mason, H. G.; Bush, J.; Agrawal, N.; Hakami, R. M.; Veneziano, R. A Microfluidic Platform to Monitor Real-Time Effects of Extracellular Vesicle Exchange between Co-Cultured Cells across Selectively Permeable Barriers. Int J Mol Sci 2022, 23 (7). DOI: 10.3390/ijms23073534 From NLM.

(52) Willms, E.; Cabañas, C.; Mäger, I.; Wood, M. J. A.; Vader, P. Extracellular Vesicle Heterogeneity: Subpopulations, Isolation Techniques, and Diverse Functions in Cancer Progression. Frontiers in Immunology 2018, 9, Review. DOI: 10.3389/fimmu.2018.00738.

(53) Reigle, J.; Secic, D.; Biesiada, J.; Wetzel, C.; Shamsaei, B.; Chu, J.; Zang, Y.; Zhang, X.; Talbot, N. J.; Bischoff, M. E.;, et al. Tobacco smoking induces metabolic reprogramming of renal cell carcinoma. J Clin Invest 2021, 131 (1). DOI: 10.1172/JCI140522 From NLM Medline.

(54) Qiu, K.; Zou, W.; Fang, Z.; Wang, Y.; Bell, S.; Zhang, X.; Tian, Z.; Xu, X.; Ji, B.; Li, D.;, et al. 2D MoS(2) and BN Nanosheets Damage Mitochondria through Membrane Penetration. ACS Nano 2023, 17 (5), 4716–4728. DOI: 10.1021/acsnano.2c11003 From NLM Medline.

(55) FastQC: A quality control tool for high throughput sequence data. http://www.bioinformatics.babraham.ac.uk/projects/fastqc (accessed 2/8/24).

(56) A wrapper tool around Cutadapt and FastQC to consistently apply quality and adapter trimming to FastQ files, with some extra functionality for MspI-digested RRBS-type (Reduced Representation Bisufite-Seq) libraries. https://www.bioinformatics.babraham.ac.uk/projects/trim_galore (accessed 2/8/24).

(57) Martin, M. Cutadapt removes adapter sequences from high-throughput sequencing reads. 2011 2011, 17 (1), 3, next generation sequencing; small RNA; microRNA; adapter removal. DOI: 10.14806/ej.17.1.200.

(58) Frankish, A.; Carbonell-Sala, S.; Diekhans, M.; Jungreis, I.; Loveland, J. E.; Mudge, J. M.; Sisu, C.; Wright, J. C.; Arnan, C.; Barnes, I.;, et al. GENCODE: reference annotation for the human and mouse genomes in 2023. Nucleic Acids Res 2023, 51 (D1), D942–d949. DOI: 10.1093/nar/gkac1071 From NLM.

(59) Dobin, A.; Davis, C. A.; Schlesinger, F.; Drenkow, J.; Zaleski, C.; Jha, S.; Batut, P.; Chaisson, M.; Gingeras, T. R. STAR: ultrafast universal RNA-seq aligner. Bioinformatics 2012, 29 (1), 15–21. DOI: 10.1093/bioinformatics/bts635 (acccessed 2/8/2024).

(60) Love, M. I.; Huber, W.; Anders, S. Moderated estimation of fold change and dispersion for RNA-seq data with DESeq2. Genome Biology 2014, 15 (12), 550. DOI: 10.1186/s13059-014-0550-8.

(61) Rozowsky, J.; Kitchen, R. R.; Park, J. J.; Galeev, T. R.; Diao, J.; Warrell, J.; Thistlethwaite, W.; Subramanian, S. L.; Milosavljevic, A.; Gerstein, M. exceRpt: A Comprehensive Analytic Platform for Extracellular RNA Profiling. Cell Syst 2019, 8 (4), 352–357.e353. DOI: 10.1016/j.cels.2019.03.004 From NLM.

(62) O’Leary, N. A.; Wright, M. W.; Brister, J. R.; Ciufo, S.; Haddad, D.; McVeigh, R.; Rajput, B.; Robbertse, B.; Smith-White, B.; Ako-Adjei, D.;, et al. Reference sequence (RefSeq) database at NCBI: current status, taxonomic expansion, and functional annotation. Nucleic Acids Research 2015, 44 (D1), D733–D745. DOI: 10.1093/nar/gkv1189 (acccessed 2/8/2024).

(63) Subramanian, A.; Tamayo, P.; Mootha, V. K.; Mukherjee, S.; Ebert, B. L.; Gillette, M. A.; Paulovich, A.; Pomeroy, S. L.; Golub, T. R.; Lander, E. S.;, et al. Gene set enrichment analysis: A knowledge-based approach for interpreting genome-wide expression profiles. Proceedings of the National Academy of Sciences 2005, 102 (43), 15545–15550. DOI: doi:10.1073/pnas.0506580102.

(64) Langevin, S. M.; Kuhnell, D.; Biesiada, J.; Zhang, X.; Medvedovic, M.; Talaska, G. G.; Burns, K. A.; Kasper, S. Comparability of the small RNA secretome across human biofluids concomitantly collected from healthy adults. PLoS One 2020, 15 (4), e0229976. DOI: 10.1371/journal.pone.0229976 From NLM Medline.

(65) Walsh, K. B.; Zimmerman, K. D.; Zhang, X.; Demel, S. L.; Luo, Y.; Langefeld, C. D.; Wohleb, E.; Schulert, G.; Woo, D.; Adeoye, O. miR-181a Mediates Inflammatory Gene Expression After Intracerebral Hemorrhage: An Integrated Analysis of miRNA-seq and mRNA-seq in a Swine ICH Model. J Mol Neurosci 2021, 71 (9), 1802–1814. DOI: 10.1007/s12031-021-01815-9 From NLM Medline.

